# Deep Neural Networks Explain Spiking Activity in Auditory Cortex

**DOI:** 10.1101/2024.11.12.623280

**Authors:** Bilal Ahmed, Joshua D. Downer, Brian J. Malone, Joseph G. Makin

## Abstract

For static stimuli or at gross (*∼*1-s) time scales, artificial neural networks (ANNs) that have been trained on challenging engineering tasks, like image classification and automatic speech recognition, are now the best predictors of neural responses in primate visual and auditory cortex. It is, however, unknown whether this success can be extended to spiking activity at fine time scales, which are particularly relevant to audition. Here we address this question with ANNs trained on speech audio, and acute multi-electrode recordings from the auditory cortex of squirrel monkeys. We show that layers of trained ANNs can predict the spike counts of neurons responding to speech audio and to monkey vocalizations at bin widths of 50 ms and below. For some neurons, the ANNs explain close to all of the explain able variance—much more than traditional spectrotemporal– receptive-field models, and more than untrained networks. Non-primary neurons tend to be more predictable by deeper layers of the ANNs, but there is much variation by neuron, which would be invisible to coarser recording modalities.

## Introduction

The primate auditory system transforms incoming acoustic information dispersed in time and frequency into distinct auditory “objects” that can be interpreted, localized, and integrated with information from the other senses. A human listener can resolve, for example, a monophonic musical recording into piano and guitar, or again into more abstract “objects” like a melody over a sequence of chords. Although two decades of electrophysiology have yielded precise characterizations of many of the tuning properties of neurons in auditory cortex, we currently do not understand how these together underwrite such computations.

On the other hand, we now have in deep artificial neural networks (ANNs) a more easily investigated system that—as of this last decade—solves such problems at human performance levels. Although crude as biophysical models, ANNs strongly resemble biological neural networks in terms of *computation* and *representation* (1–5). Recently, this has been demonstrated convincingly for the ventral visual pathway (1, 3, 4): the deep layers of ANNs trained as image classifiers explain more than half the variance of single-unit responses in areas V4 and IT atop the visual hierarchy, more than any other modeling approaches (6). This is despite the fact that these ANNs were not trained to predict activity in the brain, but only to complete an ecologically relevant task (image classification); the connection between the artificial and real neural activity was made only by way of a few hundred parameters of a linear map.

In addition to visual cortex, ANNs have been studied as encoding models for auditory cortex, as in the present study, chiefly for fMRI data (2, 7–12). The major findings from these studies are threefold: (1) ANNs trained on challenging audio “pretext” tasks (like automatic speech recognition) predict the BOLD signal in auditory cortex better than the spectrotemporal filters in classical models; (2) there is some correspondence between the hierarchical organizations of the ANNs and of auditory cortex; and (3) model prediction quality correlates with performance on the pretext task, but more strongly for some pretext tasks than others. For example, models trained to recognize speech in noise are better predicters of the BOLD signal than those trained to recognize clean speech; and models trained on multiple pretext tasks are better predictors than those trained on just one (12).

The advantage of fMRI over invasive techniques is that it allows for human subjects. On the other hand, fMRI is necessarily limited to predicting the average cortical activation (voxel intensity) over the course of the entire two-second stimulus, since it cannot resolve temporal fluctuations faster than about 1 Hz. Therefore although the studies just cited employ ANNs operating on fine time scales, their outputs are simply averaged across approximately 1 second before making predictions. This limitation is likely to be particularly destructive for the auditory system and its stimuli, which are information-rich in precisely the temporal (as opposed to spatial) dimension. Population-based decoding in core auditory cortex, for example, is optimal at a temporal resolution of less than 2 ms for discriminating sounds based on their temporal features (13). Very recent work from the Chang lab has examined ANNs as models for electrocorticography (ECoG) in the auditory cortex of humans listening to speech (14). Here, likewise, something like the three points adduced above is shown for a few different ANN architectures. For example, training ANNs on Mandarin rather than English yields better predictions of neural responses to Mandarin—in native speakers of Mandarin, but not of English. Model predictions of the ECoG envelope in the high-*γ* range were made at a fine time scale, effectively up to about 20 Hz (beyond which the analytic amplitude has no power).

What remains unclear is whether the correspondence between ANNs and auditory cortex continues to hold at the level of spiking activity. Individual ECoG channels, like fMRI voxels, represent the aggregate activity of *∼*105 neurons, so as far as these results go, the correspondence might emerge only at the population level. To explain the tuning properties of uaditory cortical neurons, it is necessary to record spiking activity.

Accordingly, we made multielectrode recordings from single- and multi-units in the core, belt, and parabelt areas of the auditory cortex of squirrel monkeys, during which the animals were exposed to a battery of *∼*600 spoken English sentences and *∼*450 monkey vocalizations. We then compared these to the responses (to the same battery of stimuli) of units in ANNs that have been trained on speech data—either fully supervised (mapping audio to letters or characters) (15–17), “weakly” supervised (18), or with a combination of supervised and unsupervised learning (19)—and with a range of different architectures and model sizes: fully convolutional, recurrent, and self-attentional. In particular, we asked how well sequences of spike counts in small (*∼*50-ms) bins can be (linearly) predicted by different layers of the ANNs. For the best network, the median correlation between model predictions and cortical responses for held-out data exceeds *∼*0.6 (after correcting for unpredictable trial-to-trial variation and removing unpredictable neurons); typically increases with layer until approximately midway through the network, although this varies by neuron and by network architecture; and exceeds that of untrained ANNs and of classical models based on spectrotemporal receptive fields (STRFs) (20). We also find some correspondence between the hierarchies of our ANNs and the traditional primary (core) vs. non-primary (belt, parabelt) distinction of primate auditory cortex.

## Results

The basic element of our analyses is the thresholded voltage on a single channel. These are likely to include multiple neurons, but for brevity and to avoid confusion with the units of the ANNs, we refer to each henceforth as a “neuron.” (We discuss spike sorting in **Methods**.)

We begin, then, with the response of such a “neuron” in monkey B to ten repetitions of a single sentence, whose spectrogram is shown on the top left in Fig. 1A. (No instance of this sentence was used in fitting the linear map from ANN to neural response; see **Methods**). In Fig. 1A (bottom left), the sequence of spike counts (in 50-ms bins) predicted by layer 2 of the Whisper [base] model is superimposed in green on top of the sequences of actual spike counts from these ten trials (gray) and their mean (black). The location of the electrode in core auditory cortex that recorded this activity is shown in green in Fig. 1B. The right half of Fig. 1B shows similar results for a recording site in a different animal (colored yellow in Fig. 1B), in response to a different sentence, and predicted by a different network (wav2vec2, layer 8). In both cases, the match is evidently quite close even at fine time scales.

**Figure 1:**
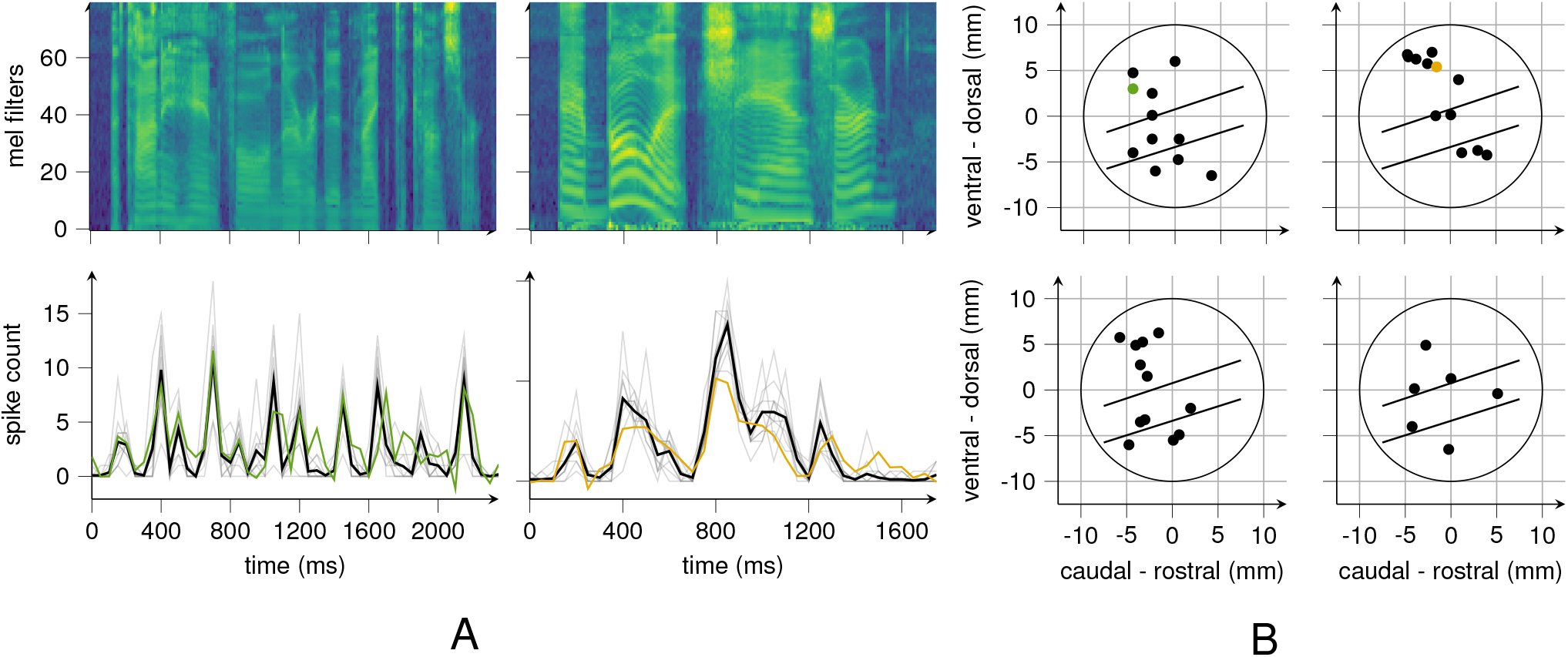
Example cortical responses and ANN predictions. (A) Left: Sequences of spike counts (50-ms bins) in response to an English sentence (“A tiny handful never did make the concert.”; spectrogram above). Shown are cortical responses to ten repetitions (gray) and their mean (black), and the prediction from the Whisper [base] ANN (green). Right: the same as the left panel but for a different sentence (“A bullet, she answered”), ANN (wav2vec2, in yellow), and monkey. The locations of the recordings are indicated by color in (B). (B) Locations of recording sites across monkeys B, C, and F. All recordings in right hemisphere except upper left panel. The large circle indicates the location of the recording cylinder, within which the upper half plane corresponds roughly to primary auditory cortex (core); the lower half, non-primary (belt and parabelt).

### Performance of ANNs as models of auditory cortex

We expand our view to all six ANN architectures, all stimuli, and the entire set of tuned neurons. We identified a neuron as tuned if its responses to multiple tokens of the same stimulus correlate with each other above chance (see **Methods**). Note that this determination is sensitive to the width of the bin in which spikes are counted; unless stated otherwise, we used 50-ms bins. About 42% (725/1718) of neurons were tuned to layer no. speech under this criterion, and 47% tuned to monkey vocalizations (see Table S1); this validates the use of speech audio as a stimulus.

To make predictions, we fit a separate linear temporal receptive field (TRF) from each layer of each ANN to each neuron (across all electrode channels, all recording sites, and all three monkeys), i.e., to the sequence of spike counts in 50-ms bins. As a baseline, we also fit a spectrotemporal receptive-field (STRFs) to each neuron. To evaluate these encoding models, we compute the correlation between sequences of spike counts and corresponding TRF predictions on a held-out set of stimuli. When reporting these correlations, we follow the standard practice of correcting for unexplainable trial-to-trial variability (21). For tuned but still highly variable neurons, this involves dividing by numbers much less than 1, which greatly increases the variance of distributions of correlations (inducing, e.g., correlations much greater than 1.0). Therefore in what follows we report correlations only for a subset of “highly tuned” neurons (253/1718) that exhibit lower levels of trial-to-trial variability (see **Methods** and Table S1).

Fig. 2 (colored shading and colored lines) shows the *distribution* of correlations across all neurons and at each layer of each ANN, in response to English sentences. First, we note that each ANN makes superior predictions to the STRF (median correlation shown as gray bar in leftmost plot and as a dashed line in all other plots) at most or all layers (i.e., the distribution of model-neuron correlations is significantly higher for the ANN layers; gray stars; Wilcoxon signed-rank test, *p <* 0.01). Indeed, the most predictive ANN layers correlate with the *median* neuron at about 0.65 (noise-corrected). This is comparable to the best models of auditory neurons to date (22), which are complex neural networks fit directly to the neural data, whereas our models fit only the linear readout.

**Figure 2:**
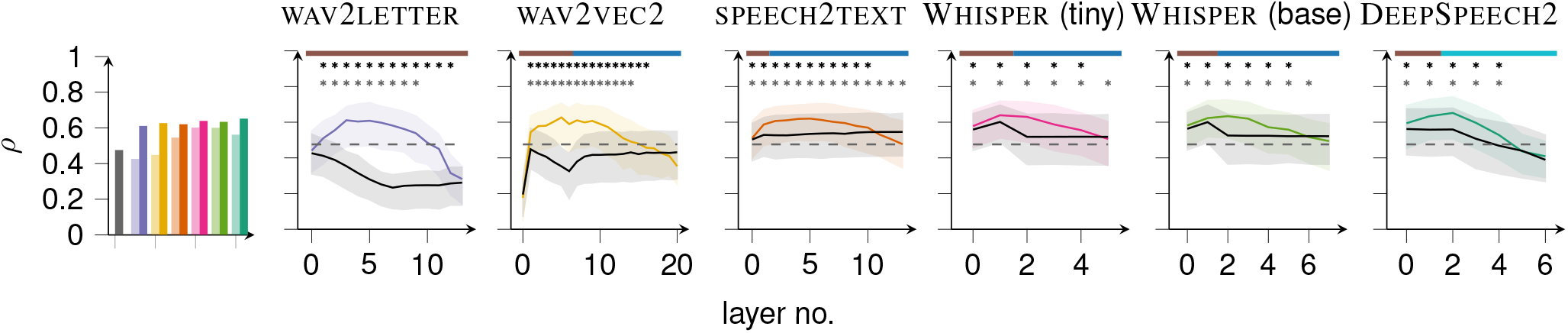
Model-neuron correlations for TIMIT stimuli. Leftmost plot: median correlation with low-variability neurons for the STRF models (gray), and for the best layers of each of the neural networks, both trained (dark colors) and untrained (light colors). Line plots: the distributions of correlations as a function of ANN layer for each of six trained networks (colored) and their untrained counterparts (gray). The median (solid line) and interquartile range (shaded region) are shown. At starred layers, the trained ANN is significantly superior to its untrained counterpart (top row of stars) or a STRF (bottom row of stars; Wilcoxon signed-rank test with *p <* 0.01). The layer type is indicated along the top of each plot: convolutional (brown), self-attention (blue), and recurrent (light blue).

Second, the distribution of correlations typically rises over the first few layers, peaks before the middle of each network, and then falls off. The rise is expected: For networks that take the raw waveform as input (wav2letter (15) and wav2vec2 (19)), it is not surprising that multiple layers of nonlinearities provide for better predictions, since the relationship between auditory-cortical responses and sound is known to be highly complex and nonlinear. But the same holds even for networks that take the spectrogram as input (speech2text (16), the Whisper models (18), and DeepSpeech2 (17)), which implies that even the STRF (which is linear in the spectrogram) is several nonlinearities away from the optimal response model. The post-peak fall off in correlations could be a result of ANN specialization for the speech-recognition task: the deepest layers predict phonemes or characters, a task that is arguably foreign to the squirrel monkey. We return to this theme in the **Discussion**.

### The importance of task optimization

To verify that task optimization improves model predictions, we investigate the predictive performance of *untrained* networks. Because each layer of an artificial neural network contains nonlinearities, a linear readout from deeper layers can be more expressive, even when the network is untrained. To determine how much of the ANN’s predictive power is due to training on the task of automatic speech recognition (ASR), and how much merely to these stacked nonlinearities, we also attempt to predict cortical activity with untrained networks. In particular, we “reset” the weights to their initial values, i.e. before training began, and then fit a new set of linear maps (TRFs) from each layer to each cortical neuron (see **Methods**). The distributions of the resulting correlations are also shown in Fig. 2, in grayscale. Trained networks are superior to untrained networks almost everywhere (Wilcoxon signed-rank test; *p <* 0.01 indicated with top row of black stars), and the most predictive layers are all in trained networks.

In networks that take the spectrogram as input, the difference beween trained and untrained networks is much smaller than in networks that take the waveform as input—presumably because in the latter, something like the spectrogram computation is learned by the trained network, but cannot be randomly assembled by the untrained network. This accords with the classical emphasis on STRFs and with the well understood transformations of the audio signal by the cochlea.

### The ecological relevance of speech recognition

In order to avail ourselves of ANNs trained with supervision on large datasets, we have let the “pretext” task for our networks be speech recognition. This in turn has narrowed our focus (up to this point) to English-sentences stimuli, since it is unclear how such networks will respond to sounds not occurring in their training sets. Three findings support this choice: (1) Roughly as many neurons—indeed, mostly the same neurons— are tuned to English and to monkey vocalizations; (2) our ANNs linearly predict the spiking responses to English sentences about as well as the best nonlinear models of auditory cortex (22); and (3) the pretext training (on speech) improves model-neuron correlations beyond what can be achieved with random nonlinearities. Although English speech is not entirely irrelevant to these animals, these results together suggest that neurons in squirrel-monkey auditory cortex are tuned to audio features that are higher-order than spectrograms but generic enough to be useful (if not used) for speech recognition, as well as processing other sounds, like monkey vocalizations. If this is the case, we expect our ANNs also to explain the responses of auditory neurons to monkey vocalizations. Fig. S1 shows the results (in the same format as Fig. 2). Note that the linear readout was re-fit for these stimuli, but the ANNs are identical to the ones analyzed in Fig. S1: they have been pretrained only on speech tasks. Nevertheless, the results are qualitatively very similar, the only notable difference being that the deepest layers of most networks tend to make better predictions of the responses to monkey vocalizations than of the responses to English speech.

Still, there is a limit to the ecological relevance of speech to the squirrel monkey. The ANNs’ performance in speech recognition does not, for example, correlate with model-neuron correlations (Fig. S2), as has been observed in studies in humans (2, 12, 14). And although in core we found about as many neurons highly tuned to speech as to monkey vocalizations, we found only about 40% as many in non-primary areas (see Table S1) (see **Discussion**).

### The time scale of predictions

Up to this point, we have performed all analyses (Figs. 1 and 2 as well as Figs. S1 and S2 and Table S1) with 50-ms bins, i.e. at 20 Hz. This represents a modeling decision: the various layers of the various ANNs operate at various sampling rates, which we have simply resampled to a single rate (20 Hz) in order to match a single bin size for counting spikes (50 ms), which is itself arbitrary. At the extreme, the biological and artificial activities could be summed over the entire stimulus-presentation period, and just one prediction made per stimulus. This is the approach that has (necessarily) been taken in the fMRI studies described previously (2, 7–12). However, the large values of correlations in Fig. 2 and the examples in Fig. 1A strongly suggest that the ANNs in this study can predict neural activity at much finer time scales than this. Here we ask more precisely what resolution yields the best correlations.

To answer this question, we low-pass filter the ANN activities with various cut-off frequencies before attempting to predict spikes binned at “high frequency” (50 Hz).

If increasing the cut-off frequency—e.g., above 10 Hz— increases model-neuron correlations, then the model is capable of explaining aspects of the neural activity that are faster than 10 Hz. Fig. 3 shows the result for a range of cut-off frequencies up to the Nyquist limit, for all six (trained) ANNs. More precisely, at each bin width, we plot the distribution of model-neuron correlations from the layer that is most predictive with 50-ms bins. The pattern is the same for all networks: model-neuron correlations increase monotonically with cutoff frequency. This shows that the ANNs are predicting fine temporal structure in the spike-count sequences, all the way up to their Nyquist limits.

**Figure 3:**
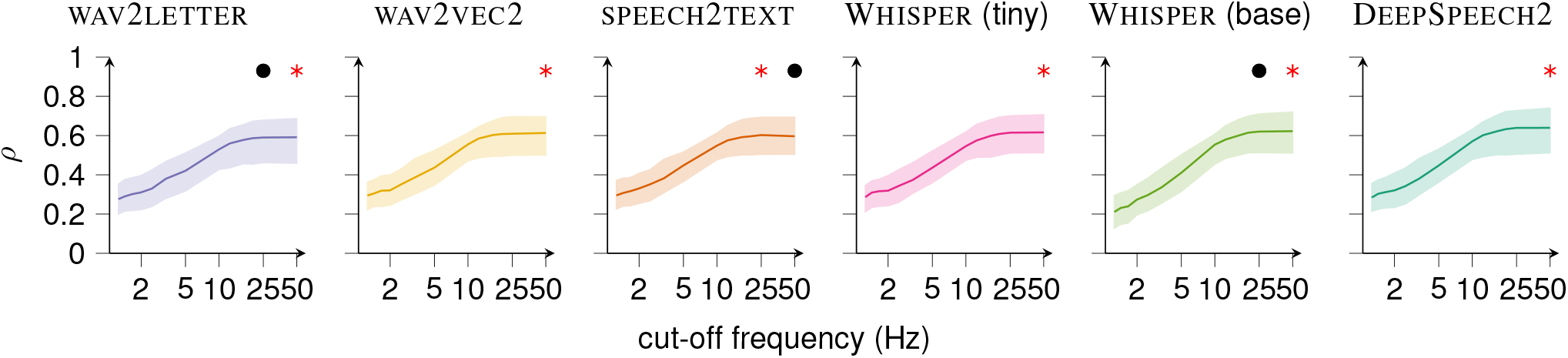
Distributions of model-neuron correlations as a function of maximum frequency of the model predictions. Median (solid line) and interquartile range (shaded) are indicated. Results are shown only for each ANN’s most predictive layer at 50-ms bins (see Fig. 2). Layer activities were low-pass filtered at the cut-off frequencies given by the horizontal axis prior to the fit of the linear readout (TRF). Position of red star corresponds to the cut-off frequency having the peak median, and black dots indicate bin widths having correlation distributions indistinguishable from that of red star (Wilcoxon signed-rank test with *p <* 0.01).

### Hierarchy

We now investigate the relationship between the hierarchies of auditory cortex and of the deep neural networks used to explain it. So far we have not distinguished between neurons recorded from core areas and those recorded in nonprimary (belt, parabelt) regions of auditory cortex. Here we ask whether a neuron’s putative location in auditory cortex (and, by implication, the auditory hierarchy) is related to the depth of the ANN layer that best predicts its responses to speech and monkey vocalizations.

For each highly tuned neuron and each ANN, we find the depth of the most predictive layer, as a fraction of the total network depth. Fig. 4 shows the distribution of these depths across all networks, and all primary (blue) and non-primary (orange) neurons. Whether the stimuli are speech or (especially) monkey vocalizations, non-primary neurons are significantly more likely than primary neurons to be best predicted by deeper layers (Wilcoxon rank-sum test, *p <* 0.05 and *p <* 0.001, resp.). The primary and non-primary distributions are broad compared to the the differences between them. This suggests a “soft” anatomical hierarchy, with many neurons in belt and parabelt playing the role of lower-level neurons, and vice versa.

**Figure 4:**
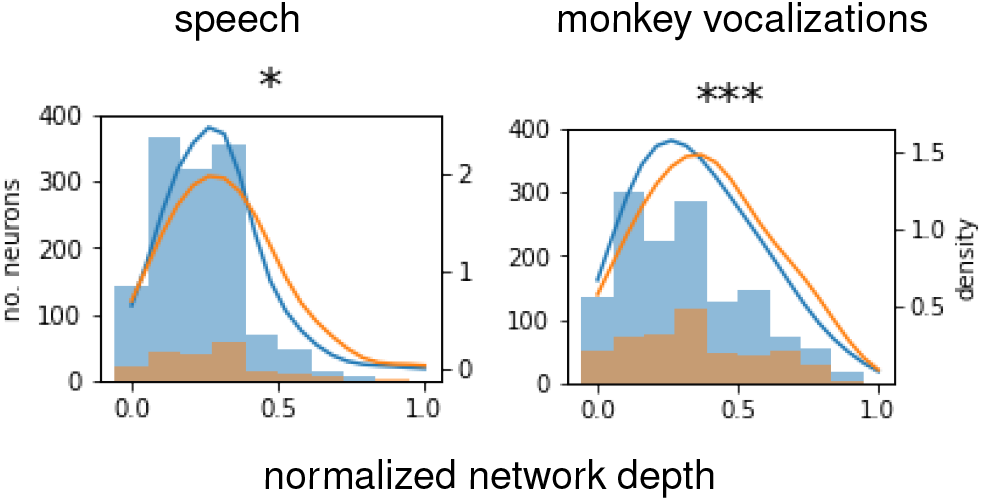
Distributions of most predictive layers (normalized) for primary (blue) and non-primary (orange) neurons. Histograms and corresponding kernel density estimates are shown as a function of network depth (from shallowest to deepest), pooled across all neurons and all six ANNs. The distribution of preferred layers is significantly “deeper” for non-primary than primary neurons (Wilcoxon rank-sum test; * for *p <* 0.05, ** for *p <* 0.01, *** for *p <* 0.001).

## Discussion

We have shown that deep neural networks make good encoding models for neurons in the auditory cortex of squirrel monkeys, even though those networks were trained, not to predict the activity of those neurons, but instead to solve a challenging auditory task, automatic speech recognition. This is (to our knowledge) the first time such an approach has been used successfully to explain spiking activity at high temporal resolution—in this case, bin widths for counting spikes as small as 20 ms.

Since the spike-count predictions are allowed to depend on a temporal *window* of layer activity (see **Methods**), each ANN based encoding model is a temporal receptive field (TRF) in the feature space of a particular ANN layer, just as the STRF is a TRF in the amplitude spectrum. The most predictive ANN layers—six or seven convolutional layers after raw waveform inputs, or about three convolutional and transformer layers after log-mel features—provide much better predictions than STRF models for most neurons. So we can say that neurons in primate auditory cortex are more tuned to the intermediate features of deep neural networks than to the amplitude spectrum. Indeed, these features explain essentially all of the explainable variance of some of the neurons from which we obtained recordings.

The present study also sheds light on the hierarchy of auditory cortex, and on the importance of task optimization, which we discuss below in connection with recent similar studies using ECoG and fMRI.

### Speech as a stimulus and speech-trained ANNs as models

We chose speech-recognition for the pretext because (1) we have found English sentences to elicit robust responses from neurons in squirrel-monkey core and belt; (2) monkey vocal calls contain somewhat similar spectrotemporal features; and (3) labeled training sets for English speech are several orders of magnitude larger than anything available for monkey calls or environmental sounds. Still, those calls do lack the complex phonemic structure of English; and, conversely, contain sounds not found in English. From this perspective, it is not surprising that the deepest layers of the ANNs do not provide as good features for explaining the activity of these neurons. Furthermore, although we found nearly the same number of core neurons tuned or highly tuned to speech as to monkey vocalizations, this ratio is much lower in non-primary areas (Table S1). This is consistent with the observation that non-primary neurons are more specialized and sparsely firing. Similarly, the explanatory power of our ANNs is on average worse for neurons in non-primary rather than in primary areas. This suggests that we are still missing important features for explaining higher-order auditory cortex.

One way to address these issues would be to train ANNs *unsupervised* on monkey vocalizations or environmental sounds. However, comparing the results of such an experiment with the results of this study is not straightforward, because it is harder to evaluate what “good” performance is on the unsupervised task. For example, if such a network outperforms the networks evaluated in this study, is it the result of better task performance or of a better task? We have therefore not taken this approach in the present study, although we consider it the clear follow up to this investigation.

### Untrained ANNs as decoding models

In our study, trained ANNs provided better encoding models than untrained ANNs; but, for ANNs that take the amplitude spectrum (i.e., the spectrogram—as opposed to a waveform) as input, untrained networks still outperform STRFs. This ordering of models held for both types of stimuli, English speech and monkey vocalizations. A recent fMRI study in humans listening to speech (9) found the same ordering and even qualitative differences among trained and untrained ANNs used as encoding models for auditory cortex: the performance of untrained networks intermediate between trained networks and STRFs, but closer to the former (see especially Fig. S3, *op. cit*.).

The result is not unexpected, although it has sometimes been obscured in the literature. Weights in ANNs are standardly initialized to small random values, which produces full-rank matrices with high probability. Furthermore, the activation functions in these ANNs are often linear (or even identity functions) near zero. Since the random weights are small and symmetrically distributed about zero, their product with inputs are also typically near zero, and therefore pass through the linear portion of the nonlinearity. So the “nonlinearities” in untrained networks are mostly information-preserving linear transformations. Consequently, the first few layers of untrained networks that take the spectrogram as input are likely to predict (linearly) neural activity as well as the spectrogram itself, i.e. as well as a STRF.

Furthermore, because the transformations are not *perfectly* linear, some new features can be created and some old features destroyed. The balance is shifted towards the former for layers that have (many) more outputs than inputs. Any such features that are (by chance) useful for predicting neural activity will be found by the linear readout that we fit to the data, and therefore provide better predictions of neural activity than the original spectrogram. This is what we observe in the first convolutional layer or two of the ANNs that take the spectrogram as input (Fig. 2, last four panels)—as expected, since the first layers have an order of magnitude more outputs than inputs.

This also explains some of the discrepancies found in the literature. Untrained networks that take the waveform, rather than the spectrogram as input, will generally provide worse models of neural activity than the STRF, because they are mostly linear in the waveform across layers. This can be seen for wav2letter and wav2vec2 in our study (Fig. 2) and for wav2vec2 in (e.g.) the recent study using ANNs as encoding models for the electrocorticogram (14). More subtly, another recent fMRI study (12) found that permuting the weights of a trained network degraded the ANNs’ performance as encoding models well below even the level of the STRF, even though those models took the spectrogram as input. But this is because training increases the size of the weights, so permutation—as opposed to re-initialization— yields large inputs to the nonlinearities, which exceed their linear regimes.

### Correspondence between the hierarchies of ANNs and auditory cortex

We found only a somewhat subtle correspondence between (on the one hand) the hierarchy of the ANNs and (on the other) the distinction between primary and non-primary auditory cortex (Fig. 4). A similar, although perhaps slightly less subtle distinction was found by Li and colleagues in ECoG data—in this case between the homologous regions in humans, Heschl’s gyrus and superior temporal gyrus, respectively (Fig. 2a, *op. cit*.). In fMRI studies (9, 12), the correspondence is more conspicuous still. Although all of these studies are in humans, rather than squirrel monkeys, the emergence of more robust hierarchical distinctions at coarser spatial and temporal granularities suggests that single units are simply more diverse in their functional role. Averaging over time and space masks this diversity.

### ANNs can predict the number of spikes in small bins

What the present study reveals, that cannot be observed from the electrocorticogram or fMRI, is that the representational correspondence with artificial neural networks emerges at the level of the single neuron, or at least single “unit” (typically 1– 5 neurons), not merely at the population level: The number of action potentials occurring in 20-ms intervals is well explained by features learned by the ANNs.

So ANNs can explain spiking activity in auditory cortex at time scales as fine as the networks’ Nyquist limits. But can they explain neural activity at time scales finer still? After all, neurons in the primate auditory cortex are known to encode information about stimuli at almost millisecond precision (23).

Here we are limited by our networks and, presumably, the pretext task. The Nyquist limits of the ANNs are set by architectural choices (e.g., stride lengths of convolutions), which could in theory be changed. However, these choices were made to achieve optimal performance on the pretext task; in particular, by the intermediate layers, sampling rates decline to about 50 Hz, which is on the order of the phoneme-production rate (10–20 Hz). Thus, complex features that require multiple layers of nonlinearities but are at finer time scales will not be predictable by such models, and the models probably cannot be changed to predict them without sacrificing performance on the pretext task.

This is another reason to explore non-speech ANNs. An interesting alternative, however, would be use *sped-up* speech as stimuli. This would retain the same enormous, labeled datasets for training; increase the *optimal* sampling rates for the ANNs; and (arguably) decrease only minimally the relevance of the stimulus to the monkey.

### Biological implausibility of networks components

Transformer attention, bidirectional recurrence, and standard convolutions are all non-causal. In theory, all of these could be remedied (with causal masking, the use of unidirectional RNNs only, and causal convolutions, respectively), although these would have adverse affects on performance on the pretext task. We have made no attempt to address these here and consider this important future work. Similarly, none of the ANNs allow information to flow from deeper to earlier layers during processing, whereas primate cortex is known to have dense feedback connections from higher to lower areas of the processing hierarchy. A new class of ANNs known as “predictive coding networks” has attempted to capture this aspect of neural computation (24); they would make potentially very interesting alternatives to standard ANNs as encoding models for the cortex.

## Methods

### Data collection

Electrophysiological signals were recorded from the auditory cortex of squirrel monkeys using a preparation described in detail elsewhere (13), which we summarize briefly here. Over the course of several months, recordings were made from penetrations into the core, belt, and parabelt areas of right auditory cortex of three animals (B, C, F). For monkey C, recordings were also made in left auditory cortex. Electrodes consisted of either 16- or a 64-channel linear probes. Auditory stimuli were played from a free-field speaker inside the soundproof chamber where recordings tooks place. All animal procedures were approved by the Institutional Animal Care and Use Committee of the University of California, San Francisco and followed the guidelines of the National Institutes of Health.

During each recording session, two sets of stimuli were presented. The first set consisted of sentences from the TIMIT corpus, each approximately 1–3 seconds in length, which we have found to elicit strong responses from the auditory cortex of squirrel monkeys. A total of 499 unique sentences were presented: 489 were presented exactly once (in random order), while the remaining 10 were repeated 11 times each, for a total of 110 presentations (also in random order). The second set consisted of monkey vocalizations, each approximately 1 second in length, including grunts, screams, and coos. In this set, a total of 303 unique vocalizations were presented: 292 were presented once, and the remaining 11 were presented 15 times each, for a total of 165 presentations, also in random order.

Spikes were identified by threshold crossing and, for the main results of this study, counted in bins of 50 ms. For Fig. 3, spikes were binned at 20 ms.

### Trial-to-trial neural variability

Neurons do not respond identically to identical stimuli, whether because they are tuned instead or additionally to other, uncontrolled stimuli; as a consequence of feedback or attention; due to false alarms or misses in spike detection; or simply because of intrinsic noise in the spiking process.

To characterize this variability, we used each neuron’s response to the repeated stimuli (see previous). In particular, since sample correlation coefficients converge on the true correlation as the number of samples (in our case, bins) increases, we first concatenated together responses from multiple trials. More precisely, we randomly selected (without replacement) a pair of responses to each of the 10 repeated sentences or each of the 11 repeated vocalizations, and concatenated together all of the first elements of each pair and likewise (in the same order) for the second elements. With 50-ms bins, this yielded two sequences of about 320 samples (20 samples/second *×* 1.6 seconds/sentence *×* 10 sentences) for speech or about 220 samples (20 samples/second *×* 1.0 second/vocalization *×* 11 vocalizations) for the vocalizations. We computed the correlation between these two sequences for each of 100,000 such random assignments, generating a distribution of trial-to-trial neural correlations for each of the *∼*1700 putative neurons (recording channels).

In order to quantify this variability, we also constructed a null distribution of correlations that would be produced by a neuron simply firing at a constant rate over the entire 320-sample (for speech) or 220-sample (for vocalizations) sequences. We computed this distribution empirically by drawing samples from a Poisson distribution with a mean firing rate of 50 spikes/s, although the rate parameter does not in fact affect the resulting distribution. The null distribution is, however, sensitive to the number of samples used to compute each correlation and therefore to the bin width. Therefore we constructed a separate null distribution for each of the stimulus sets (speech and monkey vocalizations), since their sequences lengths differed; and for the analysis in Fig. 3, which was carried out at 20-ms bins. We then put these distributions to three related but distinct uses.

#### Identifying tuned neurons

For each neuron, we asked whether its distribution of trial-to-trial response correlations is significantly greater than the null distribution under a Wilcoxon rank-sum test. With a *p*-value of 0.05, this yields 725 and 810 tuned neurons, at 50-ms bin widths, for speech and vocalizations, respectively (see Table S1).

#### Correcting for unexplainable variance

Since trial-to-trial neural variability to the same stimulus cannot be explained by any encoding model, it is common to “correct” model-neuron correlations for this excess variance. For each neuron, one first estimates the correlation between responses to identical stimuli, and then normalizes the model-neuron correlation by (a function of) this number. More precisely, there are two sources of noise in the trial-to-trial correlations (since the independent noise on the two trials) but only one in the model-neuron correlations (since the model is noise-free); so the model-neuron correlation must be normalized by the *square root* of the correlation between responses to identical stimuli. We estimated this correlation with the median of the trial-to-trial response correlations just described. This is very similar to (e.g.) the noise correction used by Pennington and David (22); they use the mean rather than median correlation.

#### Restricting to low-variability neurons

Unfortunately, distributions of correlations significantly above chance can still have very small medians. Dividing by these small numbers (or their square roots) inflates the variance in the distribution, even generating (corrected) correlations much greater than 1.0. This makes all results noisier and harder to interpret. To avoid this, we restricted our attention further to neurons whose median trial-to-trial correlation is above 90% of the probability mass of the null distribution. At 50-ms bins, there are 253 and 308 such neurons for English speech and monkey vocaliations, respectively, out of a total of 1718. This procedure is similar to that employed by Kell and colleagues (2); in their case, trial-to-trial correlations below a threshold were capped at that threshold (biasing reported correlations downward), rather than disqualifying those neurons from analysis.

### Encoding models

#### ANN architectures and pretext tasks

We considered six different neural-network architectures, all designed for speech-to-text tasks:

- wav2letter (15): 15 convolutional layers, mapping raw waveforms to letters. We modified the original network to have more slowly growing receptive fields (by reducing the convolutional kernel widths and strides), which is more computationally expensive but arguably more biologically plausible. In order to reach the same final sampling rate, we used 15 rather than 12 layers. We trained the model on 960 hours of LibriSpeech until character error rates fell below 8% on a held-out test set.
- speech2text (16), based on the Huggingface implementation (25): two convolutional layers followed by 12 transformer-encoder (self-attentional) layers, mapping log-mel filter bank to word pieces. We do not analyze the decoder. We used a model (pre)trained with standard supervised learning on LibriSpeech.
- wav2vec2 (19), based on the Huggingface implementation (26): seven convolutional layers (with layer normalization (27) and GELU activation functions (28)) followed by one convolutional layer of positional embedding, and then 12 transformer-encoder layers, mapping raw waveform to words. Our model was (pre)trained under a self-supervised contrastive loss and then only the transformer layers (and a linear projection layer following transformer layers) were fine-tuned with supervision on a speech-to-text task. The contrastive loss obliges the transformer layers to generate output that can be used to distinguish tokens masked out of its own input sequence (the output of the convolutional network) from tokens taken from other input sequences.
- DeepSpeech2 (17), based on a publicly available implementation (29): two convolutional layers followed by five bidirectional recurrent (LSTM (30)) layers, mapping log-mel filter banks to letters. We used a model (pre)trained with standard supervised learning on about 12,000 hours of read and conversational English speech (including LibriSpeech, Switchboard, WSJ, and Fisher).
- Whisper (tiny) (18) (based on the trained instance uploaded to Huggingface by OpenAI): two convolutional layers followed by four transformer-encoder layers, mapping log-mel filter banks to word pieces. We do not analyze the decoder. The model was trained to recognize speech from audio under “weak” supervision, that is, with possibly low quality labels sourced from the internet, but (consequently) upwards of 700,000 hours of audio data. About 20% of the data were non-English speech, and training also included a translation task, in addition to standard ASR.
- Whisper (base) (18) (based on the trained instance uploaded to Huggingface by OpenAI): two convolutional layers followed by six transformer-encoder layers, mapping log-mel filter bank to word pieces. We do not analyze the decoder. The model was trained the same way as Whisper (tiny).

#### Untrained ANNs

In order to distinguish the effect of training (on the pretext task) from the effects of architectural choices, we also considered the untrained counterparts to these networks. Untrained networks were obtained by (re-)initializing the network weights using the default initialization schemes for each network.

#### Predicting spiking activity with ANNs

The effective sampling rate in the ANN (imposed by the “strides” of the convolutions) declines across layers. All layer activities were therefore resampled to 20 Hz to match the 50-ms bins for the main results of this study (and for the analyses reported in Fig. 3, to 50 Hz). We then fit a temporal receptive field (TRF) from each ANN layer to each low-variability (biological) neuron (see Fig. 5). In particular, the predicted neural response at one bin was allowed to depend linearly on a 350-ms window of ANN activity. TRFs were fit on the non-repeated stimuli (see **Data collection** above) by minimizing *L*_2_-regularized squared error (*∼*20,000 samples and *∼*6000 samples at 50-ms bins for speech and monkey vocalizations, respectively). The magnitude of the *L*_2_ penalty was determined with threefold cross validation over a range of regularization parameters (10*™*5 to 1010). All fits were made with the naplib python package (31), partly customized to run on GPUs.

**Figure 5:**
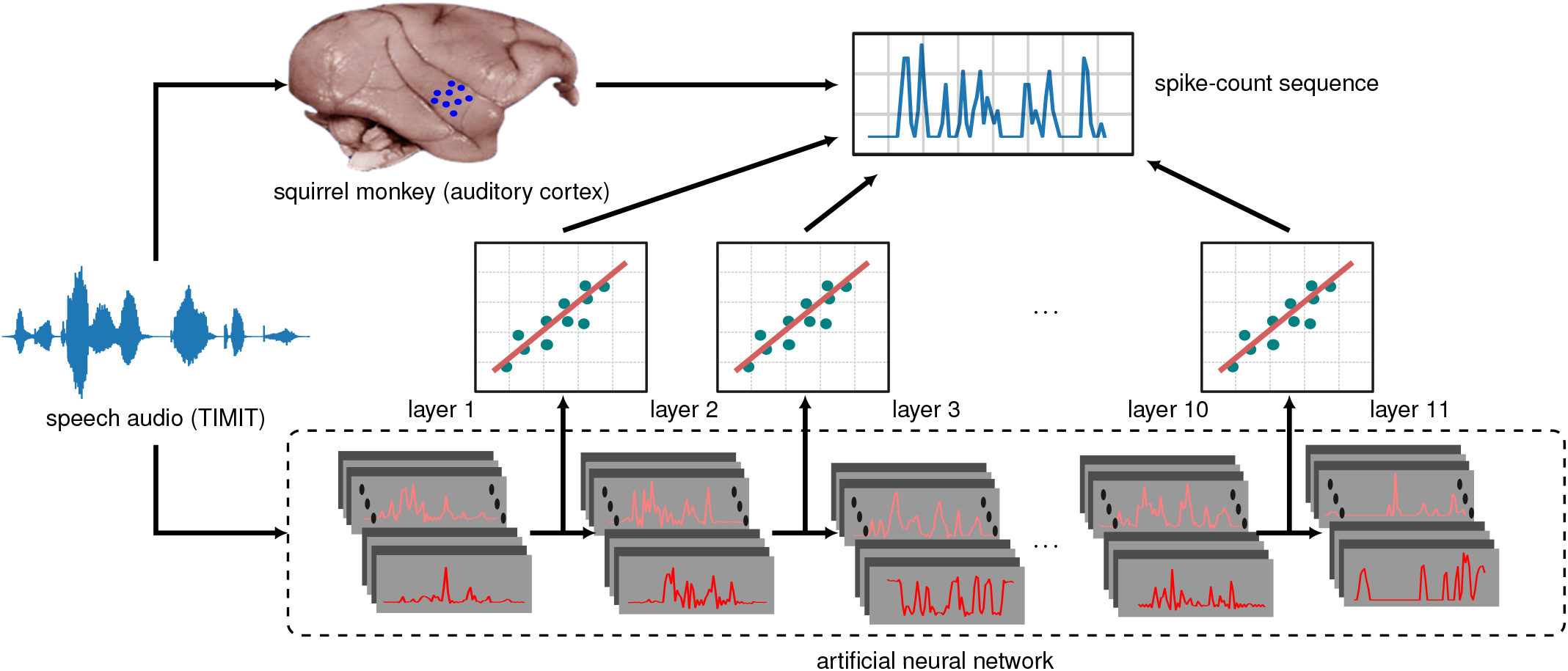
Schematic of ANN-based encoding models. Spiking activity (upper right) in response to stimuli (e.g., speech; waveform at left) is recorded from squirrel monkey auditory cortex (blue dots on brain in upper left). The exact same stimuli are presented to the ANN (bottom). The spike sequences are then regressed separately (linear plots, middle) onto each hidden layer’s responses to the same stimuli, yielding a temporal receptive field (TRF) for each layer of the ANN. Performance is evaluated on held-out stimulus-response pairs by computing the correlation between the TRF-based predictions and the spike-count sequences.

We emphasize the rationale for using a (merely) linear map: We want to be able to identify the layer of the network that best explains cortical neural activity. Allowing (arbitrary) nonlinear maps would break this correspondence, since the layers themselves are related by nonlinear maps. (We can say, colorfully, that if layer *n* makes the best predictions through a linear map, then layer 1 also makes the best predictions through the nonlinear map consisting of the neural network itself up to layer *n*, and thence out through the linear map.) At the other extreme, we are uninterested in mapping single ANN units to single neurons because a layer is arbitrary up to a (full-rank) matrix multiplication (the subsequent weight matrix could absorb the inverse of this matrix).

#### Predicting spiking activity with STRFs

As a baseline, we also fit spectrotemporal receptive fields (STRFs) (20) to every low-variability neuron. The fitting process was identical to the one just described for the ANNs, except that the input to the TRF in this case is not the ANN layer activity but the timevarying amplitude spectrum of the stimulus. For the monkey vocalizations, the amplitude spectrum was computed with a wavelet transform to mimic cochlear processing of natural sounds (31).

For the speech stimuli, however, we report STRFs fit to the log-mel filterbank computed by the input processor for speech2text (25), since we found this to yield higher model-neuron correlations than the wavelet-based computation.

### Evaluating model predictions

Our main metric of model performance is the noise-corrected correlation coefficient (see above), which we computed for every combination of low-variability neuron and encoding model. Note that an “encoding model” corresponds to a single layer from a single ANN, either trained or untrained, or to a STRF. Correlation coefficients were computed between the sequences of predicted and actual (biological) responses to the 110 and 165 held-out stimuli for speech and monkey vocalizations, respectively.

Since longer sequences yield better estimates of the correlation coefficient, we concatenated together the responses of a neuron to all of the first tokens of each stimulus type (sentence or vocalization), and likewise for the responses of the model, and computed the correlation between these long sequences. We then repeated this procedure for the second tokens of each stimulus type, and so on up through the last tokens. We report the average of these correlations coefficients across all 11 (for speech) or 15 (for vocalizations) tokens.

To compare models with each other, we compare the distributions of correlation coefficients across all low-variability neurons. (Including tuned but highly variable neurons adds noise to these comparisons, which is why we have focused on low-variability neurons.) Since the distributions to be compared consisted of paired samples (each corresponding to the same neuron), but are not Gaussian, we tested for significant differences with the Wilcoxon signed-rank test.

### Evaluating performance on the ecologically relevant pretext task

To relate the ANNs performances at prediction neural activity to their performances on the pretext task, we evaluated the word-error rates (WER) of all the six ANNs on the following datasets:

- TED-LIUM 3 (32): Test split of release 3 from the TorchAudio repository (33).
- Common Voice 5.1 (34): Test split of Common Voice 5.1 for English language (35).
- VoxPopuli (36): Test split of transcribed speech for English language from the Meta Research repository (37).

## ACKNOWLEDGEMENTS

Funding was provided by NIH 1R01DC021600-01 and the Ralph W. and Grace M. Showalter Research Trust Award. Some neural networks were trained on GPUs generously donated by the Nvidia Corporation.

## Supplementary Information

**Figure S1:**
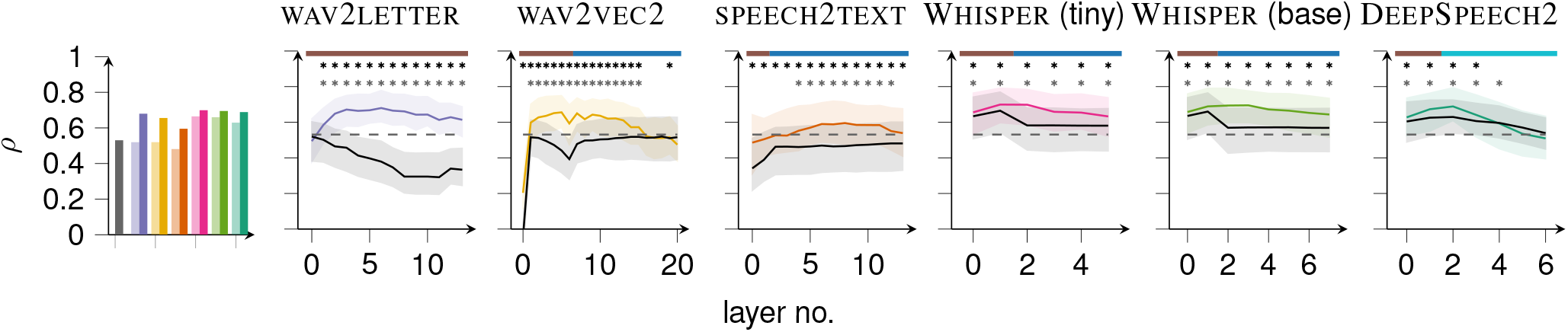
Model-neuron correlations for monkey vocalizations. Leftmost plot: median correlation with low-variability neurons for the STRF models (gray), and for the best layers of each of the neural networks, both trained (dark colors) and untrained (light colors). Line plots: the distributions of correlations as a function of ANN layer for each of six trained networks (colored) and their untrained counterparts (gray). The median (solid line) and interquartile range (shaded region) are shown. At starred layers, the trained ANN is significantly superior to its untrained counterpart (top row of stars) or a STRF (bottom row of stars; Wilcoxon signed-rank test with *p <* 0.01). The layer type is indicated along the top of each plot: convolutional (brown), self-attention (blue), and recurrent (light blue).

**Figure S2:**
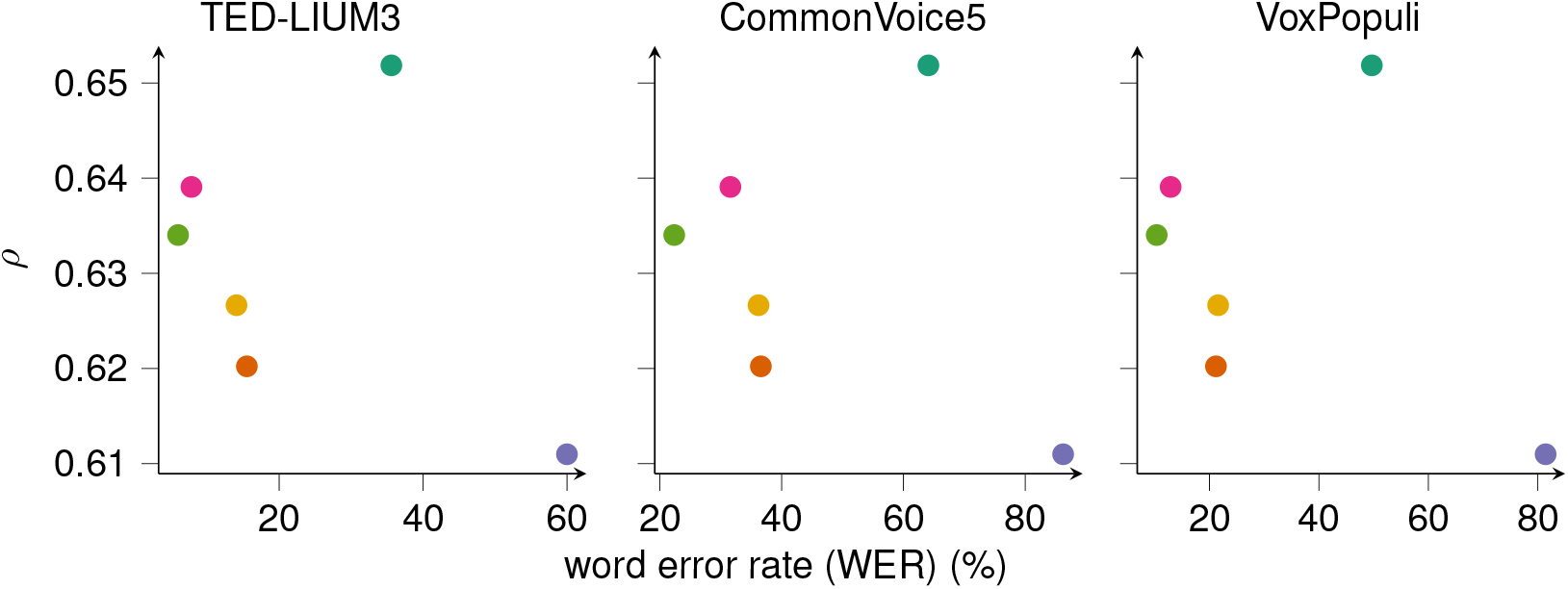
Maximum (across layers) median ANN-neuron correlation vs. ANN word error rate on three ASR data sets. Each point corresponds to an ANN (color scheme as throughout).

**Figure S3:**
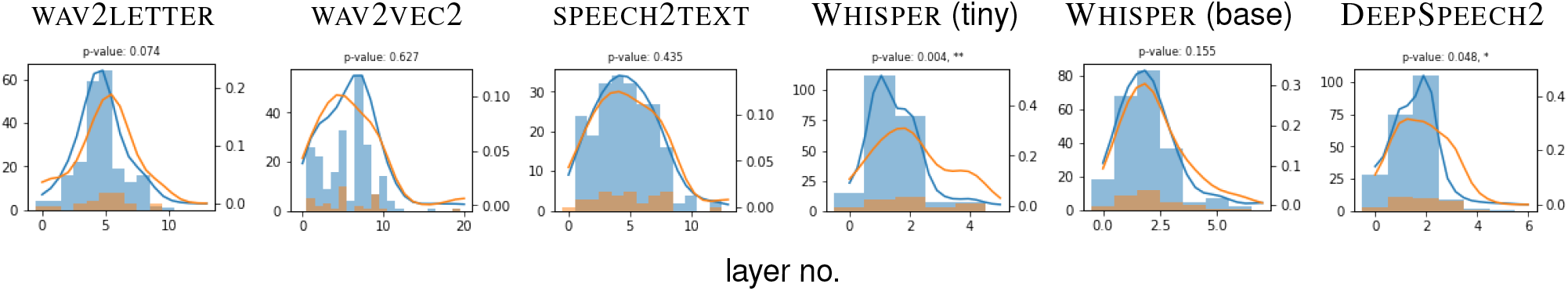
Distributions of most predictive layers (normalized) for primary (blue) and non-primary (orange) neurons, for TIMIT stimuli. Histograms and corresponding kernel density estimates are shown as a function of network depth (from shallowest to deepest) for all neurons, separately for each ANN. The distribution of preferred layers is significantly greater for non-primary than for primary neurons (Wilcoxon rank-sum test; * for *p <* 0.05, ** for *p <* 0.01, *** for *p <* 0.001).

**Figure S4:**
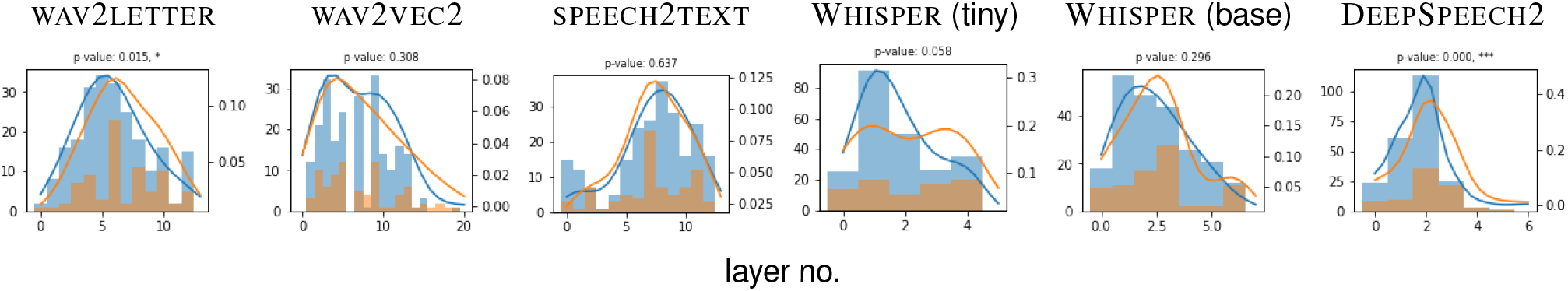
Distributions of most predictive layers (normalized) for primary (blue) and non-primary (orange) neurons, for monkey vocalizations. Histograms and corresponding kernel density estimates are shown as a function of network depth (from shallowest to deepest) for all neurons, separately for each ANN. The distribution of preferred layers is significantly greater for non-primary than for primary neurons (Wilcoxon rank-sum test; * for *p <* 0.05, ** for *p <* 0.01, *** for *p <* 0.001).

**Table S1:** **Number of tuned neurons**. Tuning is determined by the correlations between a neuron’s responses to identical stimuli on separate trials. A neuron is considered tuned or highly tuned (respectively) if the distribution of these correlations is distinguishable from (Wilcoxon rank-sum test, *p <* 0.05), or has a median above 90% of, a null distribution of correlations (see **Methods**). Speech stimuli were drawn from TIMIT; “vox” indicates monkey vocalizations.

|  | tuned |  |  | highly tuned |  |  |
| --- | --- | --- | --- | --- | --- | --- |
| neural area | speech | vox | both | speech | vox | both |
| core | 433 | 437 | 334 | 220 | 227 | 172 |
| Non-primary | 292 | 373 | 216 | 33 | 81 | 23 |
| all | 725 | 810 | 550 | 253 | 308 | 195 |

**Table S2:** **Properties of** wav2letter **(modified)**. RF: receptive field; *T*_s_: sampling period; *ρ*: median correlation.

| layer | type | RF(ms) | $T_s$(ms) | free parameters | $\rho$ |
| --- | --- | --- | --- | --- | --- |
| 0 | conv | 1.94 | 1.25 | 250 |  |
| 1 | conv | 4.44 | 2.50 | 250 |  |
| 2 | conv | 9.44 | 5.00 | 250 |  |
| 3 | conv | 19.4 | 10.0 | 250 |  |
| 4 | conv | 39.4 | 20.0 | 250 |  |
| 5 | conv | 79.4 | 20.0 | 250 |  |
| 6 | conv | 119.4 | 20.0 | 250 |  |
| 7 | conv | 159.4 | 20.0 | 250 |  |
| 8 | conv | 279.4 | 20.0 | 250 |  |
| 9 | conv | 399.4 | 20.0 | 250 |  |
| 10 | conv | 519.4 | 20.0 | 250 |  |
| 11 | conv | 639.4 | 20.0 | 250 |  |
| 12 | conv | 1239.4 | 20.0 | 2000 |  |
| 13 | conv | 1239.4 | 20.0 | 2000 |  |

**Table S3:** **Properties of** wav2vec2 RF: receptive field; *T*_s_: sampling period; *ρ*: median correlation.

| layer | type | RF(ms) | $T_s$(ms) | free parameters | $\rho$ |
| --- | --- | --- | --- | --- | --- |
| 0 | conv | 0.625 | 0.312 | 512 |  |
| 1 | conv | 1.25 | 0.625 | 512 |  |
| 2 | conv | 2.50 | 1.25 | 512 |  |
| 3 | conv | 5.00 | 2.50 | 512 |  |
| 4 | conv | 10.0 | 5.00 | 512 |  |
| 5 | conv | 15.0 | 10.0 | 512 |  |
| 6 | conv | 25.0 | 20.0 | 512 |  |
| 7-20 | attention | 2565.0 | 20.0 | 768 |  |

**Table S4:**
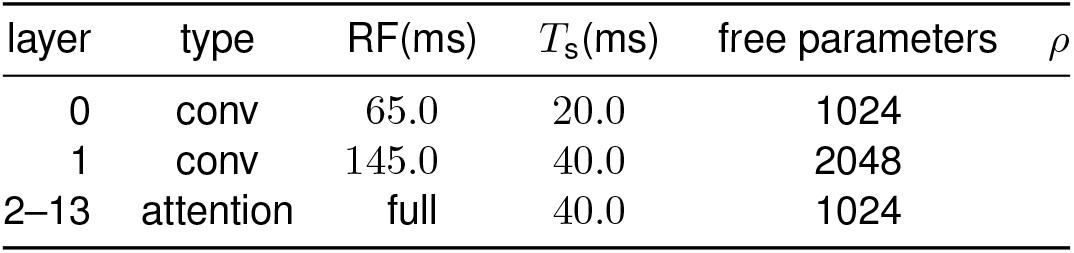
**Properties of** speech2text. RF: receptive field; *T*_s_: sampling period; *ρ*: median correlation.

**Table S5:** **Properties of** Whisper **(tiny)**. RF: receptive field; *T*_s_: sampling period; *ρ*: median correlation.

| layer | type | RF(ms) | $T_s$(ms) | free parameters | $\rho$ |
| --- | --- | --- | --- | --- | --- |
| 0 | conv | 45.0 | 10.0 | 384 |  |
| 1 | conv | 65.0 | 20.0 | 384 |  |
| 2–5 | attention | full | 20.0 | 384 |  |

**Table S6:**
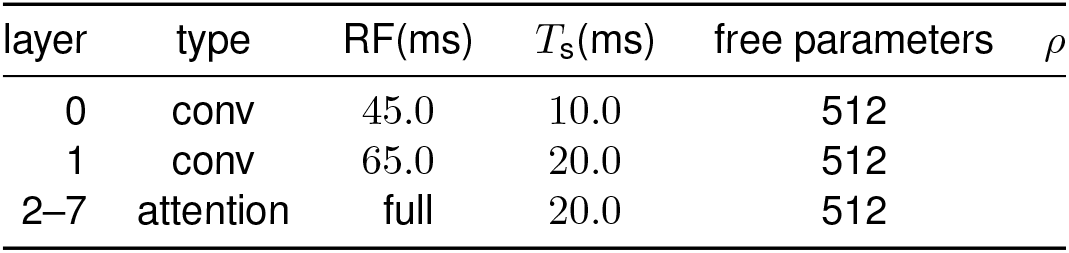
**Properties of** Whisper **(base)**. RF: receptive field; *T*_s_: sampling period; *ρ*: median correlation.

**Table S7:** **Properties of** DeepSpeech2. RF: receptive field; *T*_s_: sampling period; *ρ*: median correlation. The recurrent layers are bidirectional, LSTM-based.

| layer | type | RF(ms) | $T_s$(ms) | free parameters | $\rho$ |
| --- | --- | --- | --- | --- | --- |
| 0 | conv | 120 | 20.0 | 2592 |  |
| 1 | conv | 320 | 20.0 | 1312 |  |
| 2–6 | recurrent | full | 20.0 | 2048 |  |

